# Thyroid hormones act as a timer for the postnatal maturation of parvalbumin neurons in the neocortex

**DOI:** 10.1101/2024.12.13.628336

**Authors:** Juan Ren, Suzy Markossian, Romain Guyot, Denise Aubert, Jacques Brocard, Jiemin Wong, Frédéric Flamant, Sabine Richard

## Abstract

Thyroid hormones (THs) are required for proper maturation of parvalbumin-expressing GABAergic interneurons in the mouse neocortex, which function is essential to maintain a proper balance between neuronal excitation and inhibition. However, the timeline of this TH action has yet to be elucidated. Here, we used several mouse models expressing dominant negative mutations of the TH nuclear receptor α1 (TRα1), to better define the time window during which THs promote the maturation of parvalbumin neurons. The action of THs during the first three postnatal weeks appears to be necessary to initiate the elaboration of a specialized extracellular matrix called the perineuronal net. Transcriptome analysis identified a small set of putative target genes for TRα1, which are at the origin of the postnatal remodeling of the repertoire of ions channels and the elaboration of perineuronal nets. These data suggest that THs act as a timer to define the temporal boundaries of the critical period of heightened cortical plasticity, which plays a fundamental role in the development of neuronal circuits.

**Significance statement:** Thyroid hormones exert a major influence on late neurodevelopment, acting directly on gene expression by binding to nuclear receptors present in all cell types. This study reports the discovery of a direct link between the transcriptional regulation exerted by the thyroid hormones in GABAergic neurons and the timing of the critical period of heightened plasticity. This provides a general framework to understand the pathology associated with altered thyroid hormone signaling.

## Introduction

Parvalbumin-expressing neurons (PV neurons) are GABAergic inhibitory interneurons that play a central role in the organization of neuronal circuits in the neocortex (Hijazi, Smit et al., 2023). They are fast-spiking and synchronize network oscillations, which are thought to play an important role in information processing and memory formation. In mice, PV neurons begin to express parvalbumin during the first postnatal weeks (del Rio, de Lecea et al., 1994, Huang, Kirkwood et al., 1999). During this time, the electrical activity of these neurons progressively increases (Okaty, Miller et al., 2009) and synaptogenesis is very active. This opens the so-called critical period of heightened synaptic plasticity (Hensch, 2005) during which sensory input and experience shape neuronal networks in the neocortex. The critical period ends when circuit rewiring becomes actively dampened (Takesian & Hensch, 2013). Notably, the elaboration of the perineuronal net (PNN) around PV neurons, a densely packed extracellular matrix enriched in proteoglycans, stabilizes synapses and restricts plasticity (Wingert & Sorg, 2021). Defects in PV neuron maturation during the critical period cause a persistent imbalance between neuronal excitation and inhibition, which is suspected to be at the origin of several pathological conditions, including attention deficit hyperactivity disorder, autism, and epilepsy (Sohal & Rubenstein, 2019).

THs (including T3 or 3,3’,5-triiodo-L-thyronine, and the less active precursor T4 or thyroxine) exert a broad influence on brain development, and they are critically required for proper maturation of PV neurons (Bernal, Morte et al., 2022, Ren & Flamant, 2023, Richard, Ren et al., 2023). THs directly act on gene transcription by binding to the TRα1 and TRβ1/2 nuclear receptors, which are encoded by the *Thra* and *Thrb* genes in mice. In both humans and mice, TRα1 mutations have dramatic and irreversible consequences on neurodevelopment, and these consequences are similar to those of TH deficiency, with most human presenting with mild to severe intellectual disability (Berbel, Navarro et al., 2014, van Gucht, Moran et al., 2017).

In the postnatal mouse neocortex, PV neurons display a high sensitivity to TRα1 mutation (Wallis, Sjogren et al., 2008) and TH deficiency (Mayerl & Heuer, 2023, Uchida, Hasuoka et al., 2021). We have previously found that expressing TRα1^L400R^, a dominant-negative TRα1 receptor, only in the GABAergic lineage from an early neurodevelopmental stage, was sufficient to dramatically inhibit the expression of parvalbumin and the elaboration of PNNs (Richard, Guyot et al., 2020). By contrast, we have recently observed that expressing TRα1^L400R^ specifically in PV neurons had no visible effects on their maturation and had only moderate consequences on their function (Okaty et al., 2009). In the latter study, the specific blockade of TH signaling in PV neurons was driven by the expression of the *Pvalb* gene, which starts very gradually and with considerable temporal variation between PV neurons, ranging from a few days up to several weeks after birth (de Lecea, del Rio et al., 1995). Taken together, these results suggest that TH signaling stimulates the maturation of PV neurons during a specific time window and then becomes less important to maintain their function. Accordingly, the appearance of hypothyroidism in adult stages does not prevent the expression of parvalbumin (Gilbert, Sui et al., 2007, Uchida et al., 2021). However, the boundaries of the time window during which THs act on PV neuron maturation remain elusive. Moreover, whether blocking TH signaling during development only delays or permanently inhibits PV neuron maturation remains an open question.

To gain more insight into the timeline of TH action on PV neuron maturation, we generated novel mouse models to restrict the expression of dominant-negative TRα1 receptors in GABAergic neurons during specific time windows. Histological observations of parvalbumin and PNNs in the somatosensory cortex allowed us to pinpoint a key time window for TH action in PV neurons, around the second postnatal week. Moreover, we evidenced a lifelong persistence of defects in PV neuron maturation when TH signaling was blocked in GABAergic neurons from early stages of development. A genome-wide analysis of GABAergic neuron transcriptome in the whole neocortex two weeks after birth also allowed us to identify a set of genes whose regulation is expected to be involved in TH-mediated maturation of PV neurons. Several of these genes are notably known to be linked to the closure of the critical period. As a whole, our results show that THs, via their action on PV neurons, play a key role in cortical circuit maturation in the mouse brain.

## Results

### TRα1^E395fs401X^ expression in GABAergic neurons from an early developmental stage permanently inhibits PV neuron maturation

In our previous work, we used Cre/loxP recombination to restrict the expression of the dominant-negative TRα1^L400R^ mutant receptor in the GABAergic lineage from embryonic day 12.5 (Richard et al., 2020). As most of the mice had frequent epileptic seizures and died before postnatal day 21 (PND21), we could not address a possible recovery of PV neuron maturation during adulthood. We thus used here a similar model, based on the *Thra^Slox^* allele, encoding TRα1^E395fs401X^, a receptor with moderate dominant-negative activity (Markossian, Guyot et al., 2018). The floxed *ROSA-tdTomato^lox^*reporter transgene was also introduced to trace the cells belonging to the GABAergic lineage. One third of *Thra^Slox/+^ ROSA-tdTomato^lox/+^ Gad2-Cre* mice survived to adulthood. We therefore compared these mutant mice and *Thra^+/+^ ROSA-tdTomato^lox/+^ Gad2-Cre* littermates (controls), to histologically study whether TRα1 mutation only delays or permanently inhibits PV neuron maturation.

At PND21, while the red fluorescence did not indicate any cell loss in the GABAergic lineage (Suppl. Fig. 1), the density of parvalbumin-positive (PV+) cells was drastically lower in mutants than in controls (Fig. 1a, b). PNNs were labeled using Wisteria Floribunda Agglutinin (WFA), which binds N-acetyl-galactosamine-β1 residues of PNN glycoproteins. The density of WFA-positive (WFA+) cells was also significantly lower in mutants (Fig. 1a, c). Interestingly, the two phenotypes were accompanied by a clearly lower ratio of PV+ cells that were surrounded by PNNs in mutants, compared to controls (Fig. 1a, d). At adult stage, the same analysis showed signs of some recovery of PV neuron density in mutants (Fig. 1e, f), while the PNN defect persisted (Fig. 1e, g and h). Overall, TRα1^E395fs401X^ appears to have a similar negative influence to TRα1^L400R^ and permanently impairs the maturation of PV neurons.

**Fig. 1.**
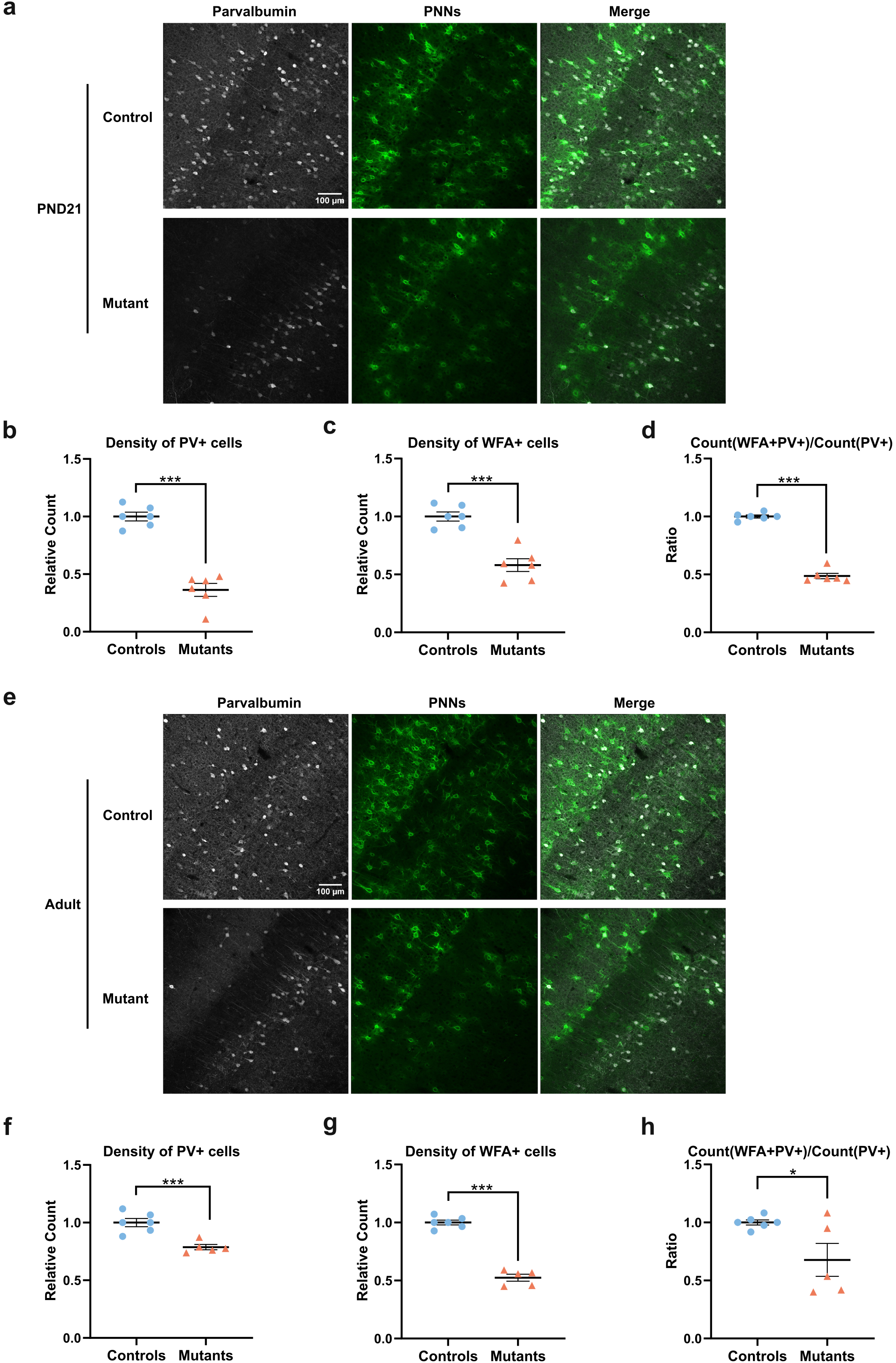
Persistent defect in PV neuron maturation in mice expressing TRα1. Histological analysis of *Thra^+/+^ ROSA-tdTomato^lox/+^ Gad2-Cre* littermates (controls) and *Thra^Slox/+^ ROSA-tdTomato^lox/+^ Gad2-Cre* mice (mutants) that express TRα1^E395fs401X^ in the GABAergic neurons and their progenitors from embryonic day 12.5 to PND21 (**a-d**, controls: n = 6, mutants: n = 6) and to adult stage (**e-h**, controls: n = 6, mutants: n = 5). **a, e** Immunohistochemistry of parvalbumin and WFA-labeled PNNs in PV neurons of somatosensory cortex at striatum level. **b, f** The density of PV+ cells was lower in mutants at both ages. **c, g** The density of WFA+ cells was lower in mutants at both ages. **d, h** The proportion of PV neurons surrounded by PNNs was lower in mutants at both ages. Data are presented as mean ± SEM and analyzed using an unpaired two-tailed T-test (\**p* < 0.05, \*\**p* < 0.01, \*\*\**p* < 0.001). Also see Suppl. Fig. 1 for related analyses.

### Postnatal expression of TRα1^L400R^ in GABAergic neurons is sufficient to inhibit PV neuron maturation

We have recently observed that the postnatal expression of TRα1^L400R^ in PV neurons has little effect on their maturation (submitted data). Although adult *Thra^AMI/+^ ROSA-tdTomato^lox/+^ Pvalb-Cre* mice displayed minor histological, electrophysiological and behavioral defects, they did not spontaneously present epileptic seizures or lethality (Ren, Markossian et al., 2024). This suggests that TH signaling intensively stimulates the maturation of PV neurons during a specific time window, and then becomes secondary to maintain their function. To better evaluate the temporal restriction of TH signaling for PV neurons maturation, we used *Thra^AMI/+^ ROSA-tdTomato^lox/+^ Gad2-CreER^T2^* mice (mutants) and *Thra^+/+^ ROSA-tdTomato^lox/+^ Gad2-CreER^T2^* or *Gad2-CreER^T2^* negative mice (controls), where Cre/loxP recombination can be induced at a chosen time by tamoxifen treatment.

A first batch of these mice were treated with tamoxifen at PND7 and histological analysis was performed at PND14. This analysis revealed that expressing TRα1^L400R^ during the second postnatal week was sufficient to compromise the onset of parvalbumin expression (Fig. 2a, b). But the amplitude of the defect in PV expression in mutant mice was not as high as that in our previous work, where TH signaling was blocked from embryonic day 12.5 to PND14 in *Thra^AMI/+^ ROSA-tdTomato^lox/+^ Gad2-Cre* mice (Richard et al., 2020). Besides, we did not observe any clear change in either the density of WFA+ cells (Fig. 2a, c) or the ratio of PV+ cells that were surrounded by PNNs (Fig. 2a, d). This may be due to the fact that the elaboration of PNNs mainly takes place after PND14, so that the density of WFA+ cells was low, with high interindividual variability. By contrast, a tamoxifen treatment performed during the fourth postnatal week (from PND21 to PND25), did not cause any significant change in either PV+ cell density or WFA+ cell density at adult stage (Fig. 2e-h). Therefore, THs exert their influence during a postnatal time window, which includes the second postnatal week and precedes weaning, since weaning usually occurs after the end of the third postnatal week in mice.

**Fig. 2.**
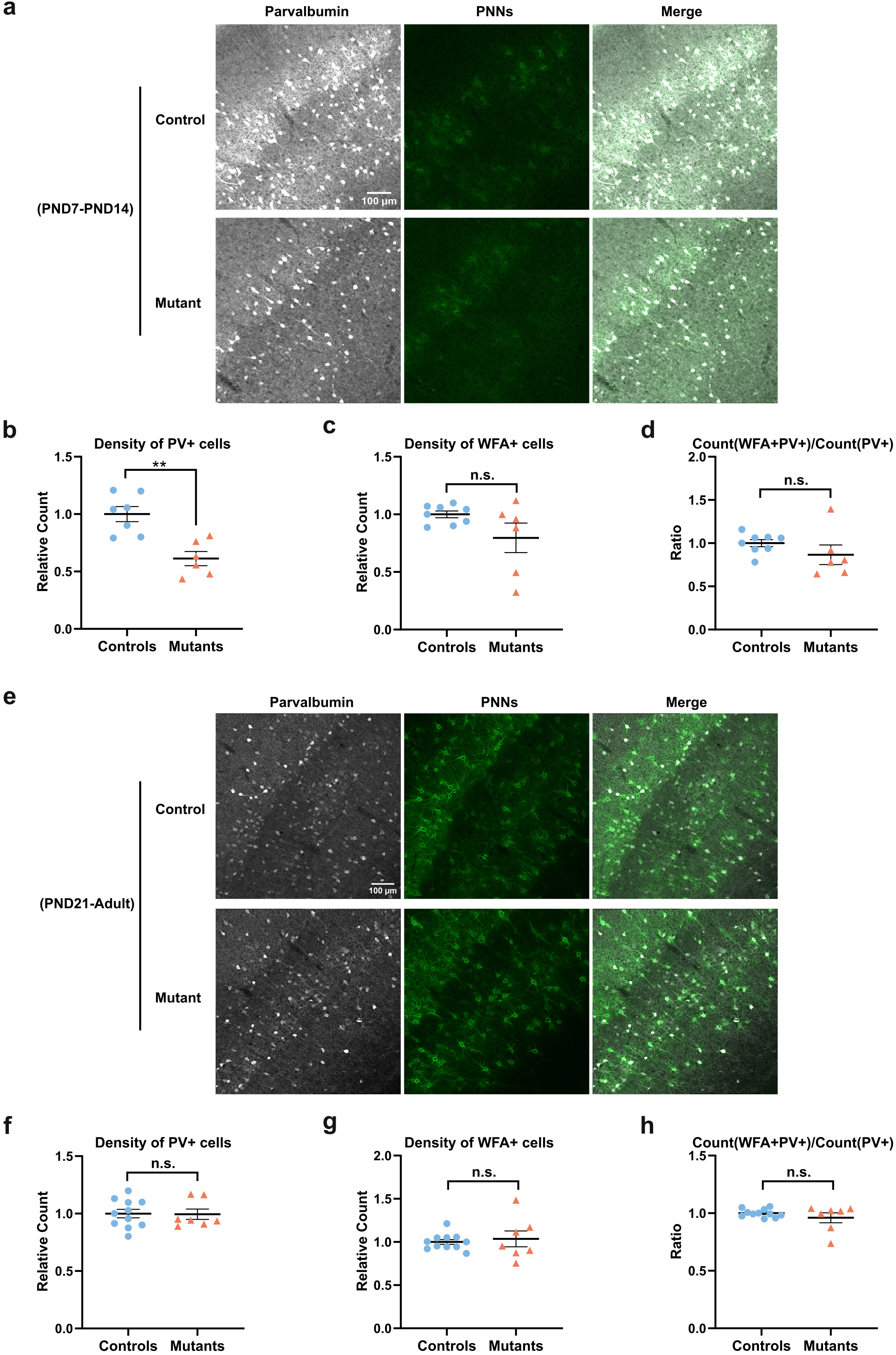
A time window for TH action on PV neuron maturation. Histological analysis of *Thra^+/+^ ROSA-tdTomato^lox/+^ Gad2-CreER^T2^* or *Gad2-CreER^T2^* negative littermates (controls) and *Thra^AMI/+^ ROSA-tdTomato^lox/+^ Gad2-CreER^T2^* mice (mutants) that express TRα1^L400R^ in the GABAergic lineage from PND7 to PND14 (**a-d**, controls: n = 8, mutants: n = 6) and from PND21 to adult (**e-h**, controls: n = 11, mutants: n = 7). **a, e** Immunohistochemistry of parvalbumin and WFA-labeled PNNs in PV neurons of somatosensory cortex at striatum level. **b, f** The density of PV+ cells was lower in mutants in former but not in latter. **c, g** The density of WFA+ cells had no clear difference in both cases. **d, h** The proportion of PV neurons surrounded by PNNs had no clear difference in both cases. Data are presented as mean ± SEM and analyzed using unpaired two-tailed T-test (\**p* < 0.05, \*\**p* < 0.01, \*\*\**p* < 0.001).

### Consequences of TRα1^L400R^ expression on gene expression in neocortical GABAergic neurons two weeks after birth

We set out to analyze the influence of TRα1^L400R^ expression on the nuclear transcriptome of GABAergic neurons at PND14, a developmental stage which appears to be quite central in the timeline of TH action in PV neurons. To do this, we used the *ROSA-GSL10gfp^lox^* reporter transgene, which labels cell nuclei, to extract nuclear RNA from neocortical GABAergic neurons, in which PV neurons represent around 50% of the population (Tasic, Menon et al., 2016). RNA-seq analysis was performed for *Thra^AMI/+^ ROSA-GSL10gfp^lox/+^ Gad2-Cre* mice and *Thra^+/+^ ROSA-GSL10gfp^lox/+^ Gad2-Cre* control littermates at PND14 Differential analysis (Deseq2 adjusted p-value > 0.05; fold-change > 2.0; basemean > 30) identified 275 down-regulated genes and 265 up-regulated genes in mutant mice (Suppl. Table 1). This large gene set contains TRα1 target genes whose transcription is normally up-regulated by liganded TRα1. Notably, among these were genes (*Hr, Klf9, Dbp, Shh, Gabrd*) that are known to be TRα1 target genes in other cell types (Zekri, Guyot et al., 2022). However, many of the observed changes in mRNA level probably represent indirect consequences of the long-term influence of TRα1^L400R^ expression on the cell physiology.

To distinguish direct TRα1 target genes from other genes that are downstream in the cascade, we carried out a series of additional studies. First, we looked for genes which are down-regulated by hypothyroidism and up-regulated by a short TH treatment at PND14. To do this, we used a propylthiouracyl (PTU) treatment to induce hypothyroidism in gestating females and their offspring. Half of the hypothyroid pups with a *Thra^+/+^ ROSA-GSL10gfp^lox/+^ Gad2-Cre* genotype then received THs at PND14. The nuclear RNA transcriptome of GABAergic neurons in the neocortex from hypothyroid pups, and from pups treated with THs for 24 hours, was analyzed at PND14 as above. In this setting we expected a selective recovery of mRNA level for TRα1 target genes.

A total of 1306 genes were sensitive to either TRα1^L400R^ expression, hypothyroidism, or TH stimulation (Fig. 3a, b and Suppl. Table 1). Among them, a large set of genes were only deregulated in hypothyroid mice. This suggests an important influence of hypothyroidism on the microenvironment (notably glial cells) of GABAergic neurons, with indirect consequences on gene expression in GABAergic neurons. The genes whose expression in GABAergic neurons was sensitive to both TRα1^L400R^ expression and hypothyroidism rather reflect the cell-autonomous influence of TH signaling in these neurons. Importantly, among a total of 388 of such genes was the *Pvalb* gene, which encodes parvalbumin. Recovery after TH treatment (TH response reached a two-fold threshold) was achieved for only a small subset of these genes (61 genes up-regulated after TH treatment and 5 genes down-regulated after TH treatment). However, clustering analysis revealed a clear trend toward recovery, which was more visible for genes up-regulated by THs (Fig. 3c) than for genes down-regulated by THs (Fig. 3d). The five genes which were down-regulated after TH treatment could reflect a poorly understood negative regulation of transcription by the liganded TRα1 (Fig. 3b). In the end, only 61 genes behaved as *bona fide* TRα1 target genes, with a robust negative response to both TRα1^L400R^ expression and hypothyroidism, and a clear induction after the short TH treatment of hypothyroid mice (Fig. 3a).

**Fig. 3.**
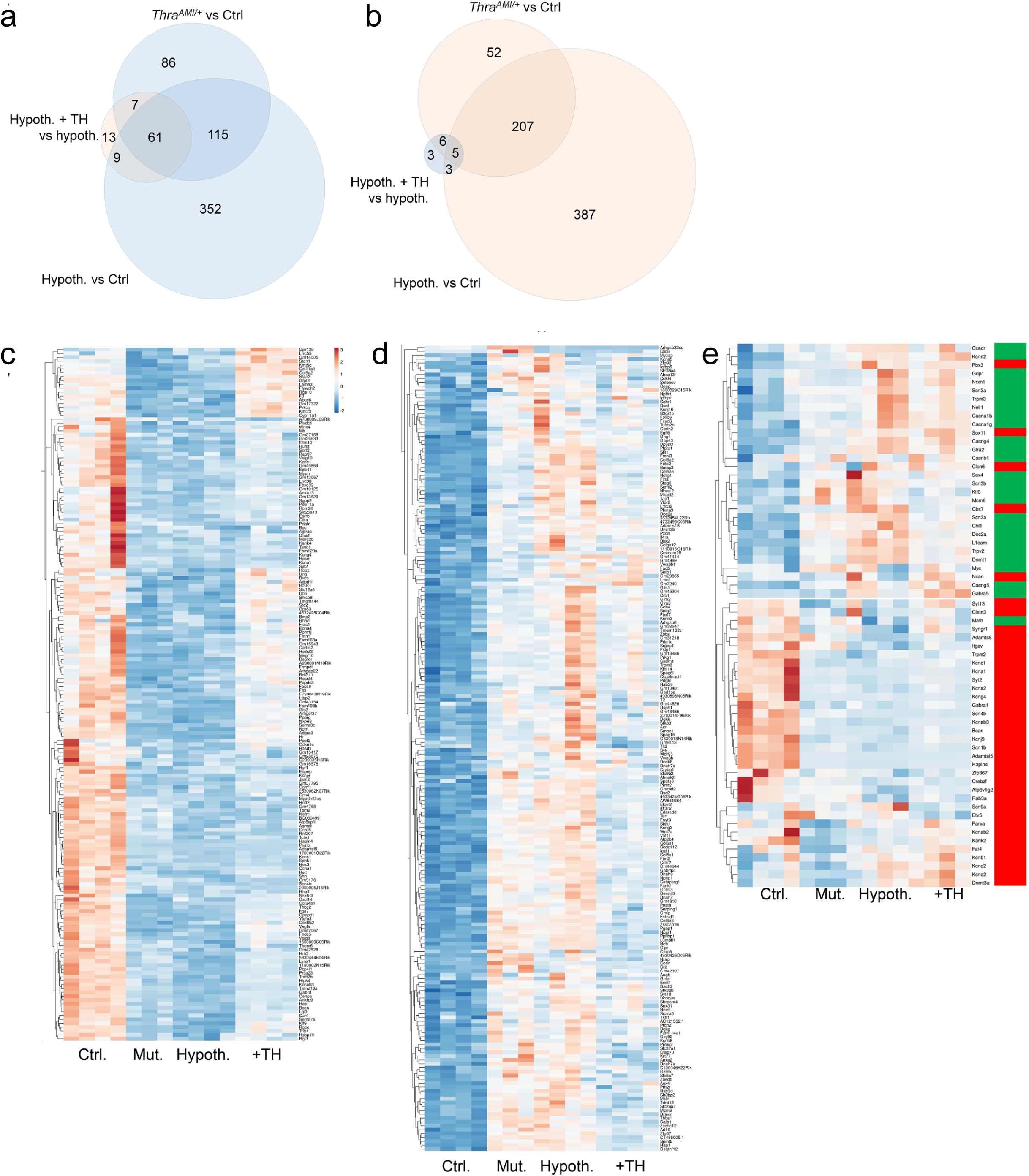
RNA-seq analysis of gene expression in sorted GABAergic neuron nuclei. Nuclei were sorted at PND14 from the neocortex of *Thra^+/+^ ROSA-GSL10gfp^lox^ Gad2-Cre* mice (Ctrl), *Thra^AMI/+^ ROSA-GSL10gfp^lox^ Gad2-Cre* mice expressing TRα1^L400R^ in GABAergic neurons (Mut), hypothyroid *Thra^+/+^ ROSA-GSL10gfp^lox^ Gad2-Cre* mice (Hypoth.), or hypothyroid *Thra^+/+^ ROSA-GSL10gfp^lox^ Gad2-Cre* mice that received a single TH injection at PND13 (+TH). **a** Venn diagram of genes that were either downregulated by TRα1^L400R^, downregulated by hypothyroidism or induced after TH treatment. **b** Venn diagram of genes that were either upregulated by TRα1^L400R^, upregulated by hypothyroidism or repressed after TH treatment (thresholds: Fold change > 2 or < 0.5, maximal adjusted base counts > 30, adjusted p-value > 0.05). **c** Heatmap showing the genes that were downregulated by both TRα1^L400R^ and hypothyroidism. Note the variable response to TH. **d** Heatmap showing the genes that were upregulated by both TRα1^L400R^ and hypothyroidism. **e** Heatmap of markers of PV neuron maturation. The marks on the right side are for genes whose expression normally increases (red) or decreases (green) between PND7 and PND21. The alterations of gene expression in mutant mice reveals a blockade of the PV neuron maturation program, caused either by TRα1^L400R^ expression or by hypothyroidism. A 24h TH treatment two weeks after birth is not sufficient to restore the expression of these marker genes. Also see Suppl. Table 1 for related datasets.

Gene Ontology analysis indicated that many of the genes deregulated in hypothyroid and mutant mice encode components of the extracellular matrix (GO:0031012; enrichment 4.4, p-value 3.10^-13^). This includes genes encoding collagens or proteins binding to the glycosaminoglycans of PNNs (GO:0005539; enrichment 4.2, p-value 5.10^-6^). This is consistent with the hypothesis that TH/TRα1 signaling stimulates the elaboration of PNNs around PV neurons. The expression of two of the downregulated genes in this family (*Sema7a* encoding semaphorin 7A; *Hapnl4* encoding hyaluronan and proteoglycan link protein 4) was restored after a short treatment with THs, suggesting that they are TRα1 target genes.

Another prominent characteristic of deregulated genes in hypothyroid and mutant mice was the presence of genes that encode voltage-gated potassium channels (GO:0005249; enrichment 6.1, p-value 10^-6^). As these genes appeared to be sensitive to acute TH treatment, their expression is unlikely to be under the direct control of TRα1. This observation suggests that hypothyroidism or long-term expression of TRα1^L400R^ in GABAergic neurons damage the postnatal remodeling of ions channels, which is a key event for proper maturation of PV neurons in the neocortex (Okaty et al., 2009). To further evaluate this possibility, we extracted the genes that mostly encode ions channels and whose expression is modified in this process of postnatal maturation, as defined by Okaty and colleagues (Okaty et al., 2009). This analysis (Fig. 3e) revealed that this postnatal transition is impaired and that the immature status of PV neurons persists in hypothyroid and mutant mice.

## Discussion

In the present study, we extend our previous observations showing that a cell-autonomous action of TH/TRα1 in GABAergic neurons is required for the maturation of neocortical PV neurons in mice. We identified a time window that spans from birth to the end of the third postnatal week, during which the action of THs is critical for the proper maturation of PV neurons. We have also found that the defects induced in PV neuron maturation when TH signaling is blocked in GABAergic neurons from early stages of development persist up to adulthood. Finally, analysis of gene expression at the end of the second week of life pinpointed a set of genes that are directly regulated by TH/TRα1 in GABAergic neurons. Some of these genes encode ion channels and PNN components, reflecting the involvement of THs in essential steps of PV neuron maturation in the developing cortex.

Using genetically modified mouse models expressing TRα1^L400R^, which exerts dominant negative activity over TRα1, we have gathered evidence suggesting that the time window during which THs promote PV neuron maturation is in the early postnatal weeks, notably during the second postnatal week. First, blocking TH/TRα1 signaling in GABAergic neurons after PND7 is sufficient to alter the expression of parvalbumin, as evidenced at PND14. On the contrary, blocking TH/TRα1 signaling in GABAergic neurons after PND21 does not significantly alter PV neuron maturation, as assessed in adulthood. PV neurons also show near-normal maturation when the blockade of TH/TRα1 signaling is restricted to PV neurons and begins only after the onset of the expression of *Pvalb*, the gene that encodes PV (Ren et al., 2024). Furthermore, a previous whole cortex RNA-seq analysis showed that gene expression changes rapidly in wild-type mice during the second postnatal week and that this evolution is stopped when TH/TRα1 signaling is blocked in GABAergic neurons (Richard et al., 2020). Finally, our novel transcriptome analysis, which was carried out with sorted GABAergic neuron nuclei, indicates that the postnatal maturation process of PV neurons is blunted by TRα1^L400R^ expression. In particular, blocking TH/TRα1 signaling appears to compromise the expression of a number of genes involved in a well-documented transition, during which the repertoire of ion channels is extensively modified (Okaty et al., 2009). It is remarkable that the second postnatal week, during which THs seem to play a crucial role in PV neuron maturation, coincides with a peak in circulating levels of THs (Friedrichsen, Christ et al., 2003). This strongly suggests that THs represent a temporal cue to synchronize the postnatal maturation of PV neurons.

The striking influence exerted by THs on cortical PV neuron maturation in the first postnatal weeks does not exclude the possibility that THs also play a role in these neurons earlier during development, or after weaning. First, we cannot fully rule out that TRα1 has an earlier function in PV neurons, as recent observations suggest that it could act prenatally to promote the tangential migration of GABAergic neuron progenitors (da Cunha Menezes, de Abreu et al., 2024). However, there is no indication that this effect is exerted in a cell-autonomous manner. Second, a potential role of THs on PV neuron maturation after weaning is supported by observations that were previously reported in mice expressing TRα1^R384C^ in all tissues, for which a complete recovery in PV neuron density was observed in adults (Wallis et al., 2008). By contrast, in the present study we have observed little recovery in PV neuron maturation in adult mice expressing TRα1^E395fs401^ in GABAergic neurons. This discrepancy is probably explained by the residual capacity of TRα1^R384C^ to bind T3 and activate transcription (Tinnikov, Nordstrom et al., 2002), which is not the case for TRα1^E395fs401^ (Richard et al., 2020). Therefore, the late onset of TRβ1 expression in PV neurons (Hochbaum, Hulshof et al., 2024) did not appear to be sufficient to counteract the influence of the dominant negative TRα1 ^E395fs401^ mutated receptor. Thus, up to now it is only for the first post-natal weeks that there is evidence for a direct, cell-autonomous, action of THs in GABAergic neurons, impacting PV neuron maturation.

The mode of action of THs in PV neurons during the second postnatal week is clarified by RNA-seq data. In the present study, we first found that PV neurons were much more sensitive to variations in TH signaling at PND14 than in adulthood, where the same method only showed minor changes in gene expression (Ren et al., 2024). Another finding of the present study was that genes that were consistently deregulated by blocking TH/TRα1 signaling in GABAergic neurons encode extracellular matrix proteins, specific PNN components or proteins interacting with PNN glycosaminoglycans. We used rapid recovery of gene expression after TH treatment in hypothyroid mice as an indication of direct transcriptional control of gene expression by TRα1. This allowed us to identify a small subset of genes whose deregulation is more likely to be the initial set of events preventing PV neuron maturation. Among these were genes encoding transcription factors or cofactors (*Dbp, Hr, Klf9*), that were already reported to be regulated by TRα1 in other contexts (Zekri et al., 2022) and that could act as upstream regulators of other deregulated genes. Moreover, in this subset of putative TRα1 target genes, we found again genes encoding PNN components, suggesting a direct regulation of PNNs by THs. Thus, it appears that liganded TRα1 might notably activate the transcription of genes encoding collagens (*Col24a1* and *Col9a2* genes), ephrin A4 (*Epha4*), laminin alpha 3 subunit (*Lama3*), semaphorin 7A (*Sema7a*) and hyaluronan and proteoglycan link protein 4 (*Hapln4*). This later protein notably represents an essential component of the PNN (Edamatsu, Miyano et al., 2018, Nojima, Miyazaki et al., 2021, Popelar, Diaz Gomez et al., 2017). Thus, a major finding of the present study is that THs directly regulate the elaboration of PNNs, which is a prominent outcome of the late maturation process of PV neurons.

The novel data presented in this paper provide strong support for the hypothesis that TH is a major temporal signal for the opening and closing of the critical period of heightened plasticity, as previously suggested (Batista & Hensch, 2019). The critical period occurs at slightly different times in different neocortical areas, as does the onset of TH signaling (Quignodon, Legrand et al., 2004). It is characterized by a series of cellular events that play a crucial role in the elaboration of functional neuronal circuits (Hensch, 2005). The beginning of parvalbumin synthesis marks the opening of the critical period, while the elaboration of PNNs around PV neurons stabilizes synapses and decides its closure. The results presented above show that TH/TRα1 signaling directly regulate the onset of parvalbumin expression, as well as the synthesis of several PNN components. Thus, our findings suggest that both the opening and the closure of the critical period are directly stimulated by liganded TRα1. Therefore, THs appear to act in the postnatal neocortex as it does during amphibian metamorphosis, as an external cue that represents an important temporal signal. During brain development, the influence of THs is also found in glial cells, as TH signaling is particularly required for timely differentiation of oligodendrocytes and myelin formation (Durand & Raff, 2000, Picou, Fauquier et al., 2014). Of note, oligodendrocyte maturation has also been shown to play a role in PV neuron maturation and in the closure of the critical period (Kalish, Barkat et al., 2020).

Patients with early TH deficiency caused by congenital hypothyroidism (Van Vliet & Deladoey, 2014) or carrying TRα1 mutations (van Gucht et al., 2017) are currently treated with TH replacement therapy. Treatment can have a positive influence on cognitive development only if it is initiated soon after birth. This is an indication that the action of THs on the maturation of the human neocortex is restricted to an early temporal window, as we have found in mice. An alteration in the timing of the critical period is also suspected to cause autism spectrum disorders (Sohal & Rubenstein, 2019). In that regard, it is noticeable that some of the rare patients in whom *THRA* mutations have been discovered were first diagnosed for autism spectrum disorders (Kalikiri, Mamidala et al., 2017, Yuen, Thiruvahindrapuram et al., 2015). Within the framework of our hypothesis, the irreversible cognitive and neurological defects that are seen both in patients with TH-related diseases and in patients with autism spectrum disorders might be similarly regarded as consequences of a disruption of the critical period of heightened cortical plasticity. We have pinpointed THs as essential players in defining the critical period. Delving deeper into the detailed molecular events occurring in the neocortex during the critical period will be key to finding therapeutic approaches to these debilitating pathologies.

### Study limitations

The complete characterization of TRα1 target genes in PV neurons would require the definition of chromatin occupancy by the receptor, as it informs of its presence on regulatory sequences. This is not currently feasible with the small number of cells that are collected when sorting GABAergic neurons from mouse neocortex. Also, the histological methods that we have used only indirectly inform of the timing of the critical period of heightened plasticity, which can be defined in a number of different ways (Quast, Reh et al., 2023, Reh, Dias et al., 2020).

## Material and methods

### Mouse models and treatments

Experiments involving the use of live animals for the current project were approved by local ethics committees (C2EA015 and C2EA017) and subsequently authorized by the French Ministry of Research (Projects #33279-2021082516194165 and #25496-2020032811174506). Mice were bred and maintained at Plateau de Biologie Expérimentale de la Souris (SFR BioSciences Gerland - Lyon Sud, France).

The *Thra^AMI^*allele allows the expression of the dominant-negative TRα1^L400R^ after the Cre/loxP-mediated deletion of a cassette with polyadenylation signals (Quignodon, Vincent et al., 2007). *Thra^Slox^* is identical, except for the presence of a C-terminus frameshift mutation which leads to the expression of the dominant-negative TRα1^E395fs401X^. The *ROSA-tdTomato^lox^* reporter transgene (also known as Ai9, MGI Cat# 4436851, RRID: MGI: 4436851) drives the Cre-dependent expression of a red fluorescent protein, mainly localized in cytoplasm (Madisen, Zwingman et al., 2010). The *ROSA-GSL10gfp^lox^*transgene encodes the GS-EGFP-L10a protein, which is a green fluorescent protein fused to the N-terminus part of the large subunit ribosomal protein L10a, mainly localized in cell nuclei (Wu, Dellinger et al., 2025). All transgenes were in C57Bl6/J genetic background. *Gad2-Cre* and *Gad2-CreER^T2^* are knock-in alleles in which the Cre recombinase reading frame is inserted into the *Gad2* gene in order to restrict the expression of the recombinase in GABAergic neurons (Taniguchi, He et al., 2011). Recombination was induced in mice carrying the *Gad2-CreER^T2^* transgene by intraperitoneal injection of tamoxifen (T5648, Sigma, dissolved in sterile sunflower oil (10 mg/ml). PND7 mice received a single 50 µL subcutaneous injection whereas 3-week-old mice received 5 daily intraperitoneal injections (daily dose: 7.5 µL/g).

Gestating mice were fed for 2 weeks with iodine deficient food supplemented with 0.15% propyl-thio uracyl (Envigo ref TD.95125) to cause deep hypothyroidism. TH levels were restored by a single intraperitoneal injection in half of the pups (10 μg T4 + 1 μg T3 dissolved in 50 μL of phosphate buffer saline, all chemicals from Sigma Aldrich France). The other PTU treated mice were injected with 50 uL of phosphate buffer saline.

### RNA-seq analysis from sorted nuclei

Nuclei were isolated from individual cortex frozen in liquid nitrogen and nuclear RNA was extracted as previously described (Wu et al., 2025). Nuclear RNA was extracted from sorted nuclei (RNeasy Micro Kit, Qiagen ref 74004) and quantified using Tapestation4150 (Agilent). 1 ng RNA was reverse-transcribed using the SMART-Seq v4 Low Input RNA kit (Takara). cDNA was quantified and qualified using Qubit (Invitrogen) and Tapestation4150 (Agilent). Libraries were then prepared from 1 ng cDNA using the Nextera XT DNA Library Kit (Illumina). Libraries were sequenced (> 2.10^7^ reads/library) on a Nextseq500 DNA sequencer (Illumina). Reads were aligned on the mouse genome (mm10 GRCm38 release) with Bowtie2 (Galaxy Version 2.4.2 + galaxy 0) (Langmead, Trapnell et al., 2009). Count table (Suppl. Table 1) was prepared using htseq-count (Galaxy Version 0.9.1 + galaxy 1, mode union, feature type: gene) (Anders, Pyl et al., 2015). Differential gene expression analysis was performed with DEseq2 (Galaxy Version 2.1.8.3) (Love, Huber et al., 2014) (Suppl. Table 1) using the following thresholds: False Discovery Rate < 0.05; p-adjusted value < 0.05; maximum adjusted basecount > 30; fold-change > 2.0. Hierarchical clustering was performed using Euclidian distance and complete distance with Clustvis (Metsalu & Vilo, 2015). Gene Ontology analysis was performed with Gorilla (https://cbl-gorilla.cs.technion.ac.il/). Raw RNA-seq data are available at NCBI Gene Expression Omnibus (GSE290072 and GSE290207).

### Brain slice collection

Each mouse was given a lethal intraperitoneal injection (6 mL/kg) of a mixture of ketamine (33 mg/mL) and xylazine (6.7 mg/mL). The thorax was opened and each mouse was perfused with 3.7% paraformaldehyde in 0.1 M phosphate buffer at room temperature. Each brain was dissected out, immersed in fixative at 4°C for 3 h and then in phosphate buffered saline (PBS) at 4°C until sectioning. Coronal sections (50 μm) were cut with the aid of a vibrating microtome (Integraslice 7550 SPDS, Campden Instruments, Loughborough, UK), in PBS at room temperature. Brain sections were stored at -20°C in cryoprotectant (30% ethylene glycol and 20% glycerol in 10 mM low-salt PBS).

### Brain slice histology

Immunohistochemistry was performed on free-floating brain sections. For PV labeling, a mouse anti-PV primary antibody (PARV19, Sigma P3088, 1:2000), and a secondary antibody made in donkey (anti-mouse DyLight 633, ThermoFisher Scientific, 1:1000) were used. For the concurrent PNN labeling, a biotinylated Wisteria floribunda lectin (WFL/WFA, Vector Laboratories B-1355, 20 µg/mL), and Streptavidin coupled to DyLight 488 (Vector Laboratories, SA-5488, 1:1000) were used. Non-specific binding sites were blocked by incubating sections for 1 h in PBS with 1% BSA and 0.2% Triton X100. Brain sections were incubated overnight at 4°C with PV antibody and biotinylated WFA, which were diluted in PBS with 1% BSA, 0.2% Triton X-100 and 1% dimethyl sulfoxide. Sections were washed in PBS and further incubated for 15 min at room temperature with DAPI (4’,6-diamidino-2-phenylindole, 1:5000, Sigma). Incubation with the secondary antibody lasted for 3 h at room temperature. Sections were mounted in Fluoroshield^TM^ (Sigma), coverslipped and imaged using an inverted confocal microscope (Zeiss LSM 780 and Zeiss LSM 800).

### Image analysis

Image analysis was performed using ImageJ. Final images resulted from the z projection of 3 optical sections (Maximum intensity tool) that were 2 µm apart (the resulting images thus reflected the fluorescence collected over a thickness of 4 µm). PV+ WFA+ Tomato+ cells were detected from final images using the Cellpose wrapper for Fiji (Stringer, Wang et al., 2021). Numbers of mono-, double- or triple-labelled cells per slice were measured automatically from 3-4 slices per mouse. The code of the original macro may be found here: https://github.com/jbrocardplatim/MacroPV The densities of Tomato+, PV+, WFA+ cells, and their co-localizations were counted in the whole area of one image taken in the somatosensory cortex area. 3-4 images in each hemisphere were analyzed for each mouse. Quantitative data were analyzed using two-tailed unpaired T-test. Data are expressed as mean ± SEM. Data analyses were performed with Excel 2016 and GraphPad Prism software (v.8.). The level of significance was set at *p* < 0.05.

## Acknowledgements

We acknowledge the contribution of SFR Biosciences (Université Claude Bernard Lyon 1, CNRS UAR3444, Inserm US8, ENS de Lyon): Nadine Aguilera and the Plateau de Biologie Expérimentale de la Souris (ANIRA-PBES) for mouse breeding; ANIRA-CYTOMETRIE and particularly Sébastien Dussurgey for nuclei sorting; ANIRA-AGC for mouse genotyping. We thank Benjamin Gillet and Sandrine Hughes of the deep sequencing facility (PSI IGFL, Lyon). We also thank Elodie Martel for carrying out histology experiments as part of her internships and Isabelle Dusart (Institut de Biologie Paris Seine, France) for helpful discussions. We acknowledge China Scholarship Council for a scholarship to Juan REN to do her PhD project in ENS de Lyon. Research fundings were from European Union’s Horizon 2020 research and innovation program, under grant agreement No. 825753 (ERGO), Agence Nationale de la Recherche (Thyromut2 program; ANR-15-CE14-0011-01).

## References

Anders S, Pyl PT, Huber W (2015) HTSeq--a Python framework to work with high-throughput sequencing data. Bioinformatics 31: 166–9

Batista G, Hensch TK (2019) Critical Period Regulation by Thyroid Hormones: Potential Mechanisms and Sex-Specific Aspects. Front Mol Neurosci 12: 77

Berbel P, Navarro D, Roman GC (2014) An evo-devo approach to thyroid hormones in cerebral and cerebellar cortical development: etiological implications for autism. Front Endocrinol (Lausanne) 5: 146

Bernal J, Morte B, Diez D (2022) Thyroid hormone regulators in human cerebral cortex development. J Endocrinol 255: R27–R36

da Cunha Menezes E, de Abreu FF, Davis JB, Maurer SV, Roshko VC, Richardson A, Dowell J, Cassella SN, Stevens HE (2024) Effects of gestational hypothyroidism on mouse brain development: Gabaergic systems and oxidative stress. Dev Biol 515: 112–120

de Lecea L, del Rio JA, Soriano E (1995) Developmental expression of parvalbumin mRNA in the cerebral cortex and hippocampus of the rat. Brain Res Mol Brain Res 32: 1–13

del Rio JA, de Lecea L, Ferrer I, Soriano E (1994) The development of parvalbumin-immunoreactivity in the neocortex of the mouse. Brain Res Dev Brain Res 81: 247–59

Durand B, Raff M (2000) A cell-intrinsic timer that operates during oligodendrocyte development. Bioessays 22: 64–71

Edamatsu M, Miyano R, Fujikawa A, Fujii F, Hori T, Sakaba T, Oohashi T (2018) Hapln4/Bral2 is a selective regulator for formation and transmission of GABAergic synapses between Purkinje and deep cerebellar nuclei neurons. J Neurochem 147: 748–763

Friedrichsen S, Christ S, Heuer H, Schafer MK, Mansouri A, Bauer K, Visser TJ (2003) Regulation of iodothyronine deiodinases in the Pax8-/- mouse model of congenital hypothyroidism. Endocrinology 144: 777–84

Gilbert ME, Sui L, Walker MJ, Anderson W, Thomas S, Smoller SN, Schon JP, Phani S, Goodman JH (2007) Thyroid hormone insufficiency during brain development reduces parvalbumin immunoreactivity and inhibitory function in the hippocampus. Endocrinology 148: 92–102

Hensch TK (2005) Critical period plasticity in local cortical circuits. Nat Rev Neurosci 6: 877–88

Hijazi S, Smit AB, van Kesteren RE (2023) Fast-spiking parvalbumin-positive interneurons in brain physiology and Alzheimer’s disease. Mol Psychiatry 28: 4954–4967

Hochbaum DR, Hulshof L, Urke A, Wang W, Dubinsky AC, Farnsworth HC, Hakim R, Lin S, Kleinberg G, Robertson K, Park C, Solberg A, Yang Y, Baynard C, Nadaf NM, Beron CC, Girasole AE, Chantranupong L, Cortopassi MD, Prouty S et al. (2024) Thyroid hormone remodels cortex to coordinate body-wide metabolism and exploration. Cell 187: 5679–5697 e23

Huang ZJ, Kirkwood A, Pizzorusso T, Porciatti V, Morales B, Bear MF, Maffei L, Tonegawa S (1999) BDNF regulates the maturation of inhibition and the critical period of plasticity in mouse visual cortex. Cell 98: 739–55

Kalikiri MK, Mamidala MP, Rao AN, Rajesh V (2017) Analysis and functional characterization of sequence variations in ligand binding domain of thyroid hormone receptors in autism spectrum disorder (ASD) patients. Autism research : official journal of the International Society for Autism Research

Kalish BT, Barkat TR, Diel EE, Zhang EJ, Greenberg ME, Hensch TK (2020) Single-nucleus RNA sequencing of mouse auditory cortex reveals critical period triggers and brakes. Proc Natl Acad Sci U S A 117: 11744–11752

Langmead B, Trapnell C, Pop M, Salzberg SL (2009) Ultrafast and memory-efficient alignment of short DNA sequences to the human genome. Genome biology 10: R25

Love MI, Huber W, Anders S (2014) Moderated estimation of fold change and dispersion for RNA-seq data with DESeq2. Genome biology 15: 550

Madisen L, Zwingman TA, Sunkin SM, Oh SW, Zariwala HA, Gu H, Ng LL, Palmiter RD, Hawrylycz MJ, Jones AR, Lein ES, Zeng H (2010) A robust and high-throughput Cre reporting and characterization system for the whole mouse brain. Nature neuroscience 13: 133–40

Markossian S, Guyot R, Richard S, Teixeira M, Aguilera N, Bouchet M, Plateroti M, Guan W, Gauthier K, Aubert D, Flamant F (2018) CRISPR/Cas9 Editing of the Mouse Thra Gene Produces Models with Variable Resistance to Thyroid Hormone. Thyroid 28: 139–150

Mayerl S, Heuer H (2023) Thyroid hormone transporter Mct8/Oatp1c1 deficiency compromises proper oligodendrocyte maturation in the mouse CNS. Neurobiol Dis 184: 106195

Metsalu T, Vilo J (2015) ClustVis: a web tool for visualizing clustering of multivariate data using Principal Component Analysis and heatmap. Nucleic Acids Res 43: W566–70

Nojima K, Miyazaki H, Hori T, Vargova L, Oohashi T (2021) Assessment of Possible Contributions of Hyaluronan and Proteoglycan Binding Link Protein 4 to Differential Perineuronal Net Formation at the Calyx of Held. Front Cell Dev Biol 9: 730550

Okaty BW, Miller MN, Sugino K, Hempel CM, Nelson SB (2009) Transcriptional and electrophysiological maturation of neocortical fast-spiking GABAergic interneurons. The Journal of neuroscience : the official journal of the Society for Neuroscience 29: 7040–52

Picou F, Fauquier T, Chatonnet F, Richard S, Flamant F (2014) Deciphering direct and indirect influence of thyroid hormone with mouse genetics. Mol Endocrinol 28: 429–41

Popelar J, Diaz Gomez M, Lindovsky J, Rybalko N, Burianova J, Oohashi T, Syka J (2017) The absence of brain-specific link protein Bral2 in perineuronal nets hampers auditory temporal resolution and neural adaptation in mice. Physiol Res 66: 867–880

Quast KB, Reh RK, Caiati MD, Kopell N, McCarthy MM, Hensch TK (2023) Rapid synaptic and gamma rhythm signature of mouse critical period plasticity. Proc Natl Acad Sci U S A 120: e2123182120

Quignodon L, Legrand C, Allioli N, Guadano-Ferraz A, Bernal J, Samarut J, Flamant F (2004) Thyroid hormone signaling is highly heterogeneous during pre- and postnatal brain development. J Mol Endocrinol 33: 467–76

Quignodon L, Vincent S, Winter H, Samarut J, Flamant F (2007) A point mutation in the activation function 2 domain of thyroid hormone receptor alpha1 expressed after CRE-mediated recombination partially recapitulates hypothyroidism. Mol Endocrinol 21: 2350–60

Reh RK, Dias BG, Nelson CA, 3rd, Kaufer D, Werker JF, Kolb B, Levine JD, Hensch TK (2020) Critical period regulation across multiple timescales. Proc Natl Acad Sci U S A 117: 23242–23251

Ren J, Flamant F (2023) Thyroid hormone as a temporal switch in mouse development. Eur Thyroid J 12

Ren J, Markossian S, Guyot R, Aubert D, Li D, Cauli B, Riet F, Wong J, Flamant F, Richard S (2024) Thyroid hormones maintain parvalbumin neuron functions in mouse neocortex. bioRxiv: 2024.07.16.603713

Richard S, Guyot R, Rey-Millet M, Prieux M, Markossian S, Aubert D, Flamant F (2020) A Pivotal Genetic Program Controlled by Thyroid Hormone during the Maturation of GABAergic Neurons. iScience 23: 100899

Richard S, Ren J, Flamant F (2023) Thyroid hormone action during GABAergic neuron maturation: The quest for mechanisms. Front Endocrinol (Lausanne) 14: 1256877

Sohal VS, Rubenstein JLR (2019) Excitation-inhibition balance as a framework for investigating mechanisms in neuropsychiatric disorders. Mol Psychiatry 24: 1248–1257

Stringer C, Wang T, Michaelos M, Pachitariu M (2021) Cellpose: a generalist algorithm for cellular segmentation. Nat Methods 18: 100–106

Takesian AE, Hensch TK (2013) Balancing plasticity/stability across brain development. Prog Brain Res 207: 3–34

Taniguchi H, He M, Wu P, Kim S, Paik R, Sugino K, Kvitsiani D, Fu Y, Lu J, Lin Y, Miyoshi G, Shima Y, Fishell G, Nelson SB, Huang ZJ (2011) A resource of Cre driver lines for genetic targeting of GABAergic neurons in cerebral cortex. Neuron 71: 995–1013

Tasic B, Menon V, Nguyen TN, Kim TK, Jarsky T, Yao Z, Levi B, Gray LT, Sorensen SA, Dolbeare T, Bertagnolli D, Goldy J, Shapovalova N, Parry S, Lee C, Smith K, Bernard A, Madisen L, Sunkin SM, Hawrylycz M et al. (2016) Adult mouse cortical cell taxonomy revealed by single cell transcriptomics. Nature neuroscience 19: 335–46

Tinnikov A, Nordstrom K, Thoren P, Kindblom JM, Malin S, Rozell B, Adams M, Rajanayagam O, Pettersson S, Ohlsson C, Chatterjee K, Vennstrom B (2002) Retardation of post-natal development caused by a negatively acting thyroid hormone receptor alpha1. Embo J 21: 5079–87

Uchida K, Hasuoka K, Fuse T, Kobayashi K, Moriya T, Suzuki M, Katayama N, Itoi K (2021) Thyroid hormone insufficiency alters the expression of psychiatric disorder-related molecules in the hypothyroid mouse brain during the early postnatal period. Sci Rep 11: 6723

van Gucht ALM, Moran C, Meima ME, Visser WE, Chatterjee K, Visser TJ, Peeters RP (2017) Resistance to Thyroid Hormone due to Heterozygous Mutations in Thyroid Hormone Receptor Alpha. Current topics in developmental biology 125: 337–355

Van Vliet G, Deladoey J (2014) Diagnosis, treatment and outcome of congenital hypothyroidism. Endocr Dev 26: 50–9

Wallis K, Sjogren M, van Hogerlinden M, Silberberg G, Fisahn A, Nordstrom K, Larsson L, Westerblad H, Morreale de Escobar G, Shupliakov O, Vennstrom B (2008) Locomotor deficiencies and aberrant development of subtype-specific GABAergic interneurons caused by an unliganded thyroid hormone receptor alpha1. The Journal of neuroscience : the official journal of the Society for Neuroscience 28: 1904–15

Wingert JC, Sorg BA (2021) Impact of Perineuronal Nets on Electrophysiology of Parvalbumin Interneurons, Principal Neurons, and Brain Oscillations: A Review. Front Synaptic Neurosci 13: 673210

Wu S, Dellinger J, Markossian S, Dusabyinema Y, Guyot R, Hughes S, Aubert D, Fackeure M, Gauthier K, Gillet B, Jiang W, Flamant F (2025) An Atlas of Thyroid Hormone Responsive Genes in Adult Mouse Hypothalamus. Endocrinology 166

Yuen RK, Thiruvahindrapuram B, Merico D, Walker S, Tammimies K, Hoang N, Chrysler C, Nalpathamkalam T, Pellecchia G, Liu Y, Gazzellone MJ, D’Abate L, Deneault E, Howe JL, Liu RS, Thompson A, Zarrei M, Uddin M, Marshall CR, Ring RH et al. (2015) Whole-genome sequencing of quartet families with autism spectrum disorder. Nature medicine 21: 185–91

Zekri Y, Guyot R, Flamant F (2022) An Atlas of Thyroid Hormone Receptors’ Target Genes in Mouse Tissues. Int J Mol Sci 23

