## Supplementary material for "Thyroid hormones act as a timer for the postnatal maturation of parvalbumin neurons in the neocortex": Suppl figure 1

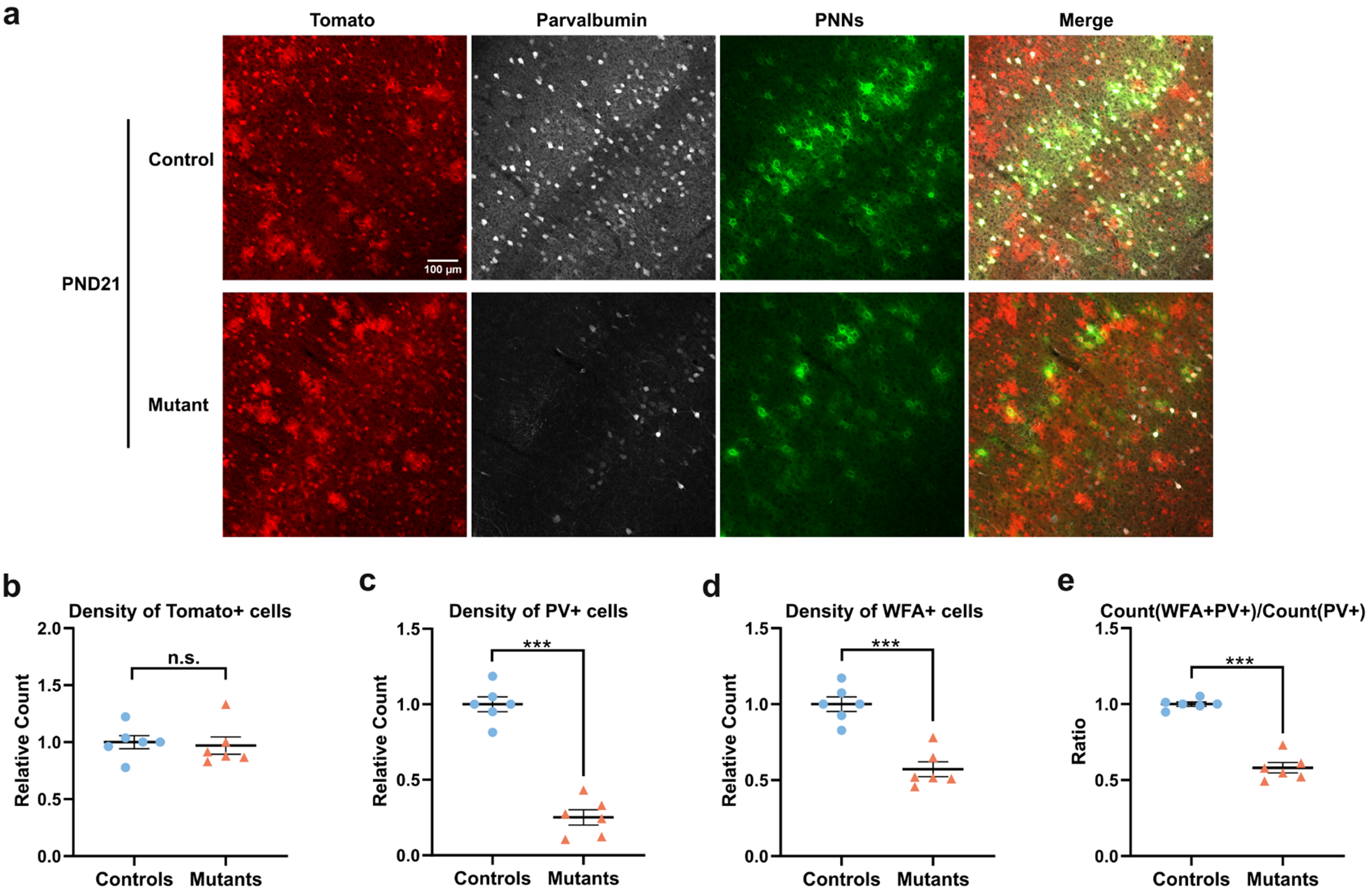


**Supplementary Fig. 1 | Persistent defect in PV neuron maturation in mice expressing TRα1$^{E395fs401X}$ in another coronal depth.** Histological analysis of *Thra$^{+/+}$ ROSA-tdTomato$^{lox/+}$ Gad2-Cre* littermates (controls) and *Thra$^{Slox/+}$ ROSA-tdTomato$^{lox/+}$ Gad2-Cre* mice (mutants) that express TRα1$^{E395fs401X}$ in the GABAergic lineage from embryonic day 12.5 to PND21 (controls: n = 6, mutants: n = 6). **a** Immunohistochemistry of parvalbumin and WFA-labeled PNNs in PV neurons of somatosensory cortex at hippocampus level. **b** The density of Tomato+ cells had no clear difference between both groups. **c** The density of PV+ cells was lower in mutants. **d** The density of WFA+ cells was lower in mutants. **e** The proportion of PV neurons surrounded by PNNs was lower in mutants. Data are presented as mean ± SEM and analyzed using unpaired two-tailed T-test (*$p < 0.05$, **$p < 0.01$, ***$p < 0.001$).
